# GeneExt: a gene model extension tool for enhanced single-cell RNA-seq analysis

**DOI:** 10.1101/2023.12.05.570120

**Authors:** Grygoriy Zolotarov, Xavier Grau-Bové, Arnau Sebé-Pedrós

## Abstract

Incomplete gene models negatively impact single-cell gene expression quantification. This is particularly true in non-model species where often gene 3′ ends are inaccurately annotated, while most scRNA-seq methods only capture the 3′ transcript region. This results in many genes being incorrectly quantified or not detected. GeneExt leverages scRNA-seq data to refine gene annotations, enhancing biological interpretation and cross-species comparisons of cell type expression atlases.

## Main

Single-cell transcriptomics has transformed the study of cell type diversity across organisms. This technology enables the large-scale and minimally biased molecular characterization of cell types at the whole-organism level, opening the window to cross-species comparisons, discovery of novel cell types, and understanding of gene regulatory programs^1^. An important problem that hampers the analysis and interpretation of scRNA-seq data in non-model species is the inaccuracy of gene annotations (missing genes, partial genes, etc.)^2–4^. The problem is aggravated by the fact that most scRNA-seq methods profile the 3′ end of the transcript, where UTRs are often particularly difficult to annotate^5–8^. Thus, a large fraction of sequencing reads map to non-genic regions of the genome and many genes are missing from single-cell expression matrices. This affects both downstream analysis (e.g. cell clustering) and the biological interpretation of the data (e.g. the possibility to miss-quantify key marker genes, or to randomly miss orthologs in cross-species comparisons^9^). Here, we introduce GeneExt, a tool that solves 3′ end annotation and other related gene annotation problems typically associated with non-model organism single-cell RNA-seq data analysis.

GeneExt takes as input scRNA-seq mapped reads and a gene annotation file (GTF or GFF, any version) and outputs extended gene annotation file ready for scRNA-seq analysis. The main functions of GeneExt are (**Fig. 1**):

**Figure 1.**
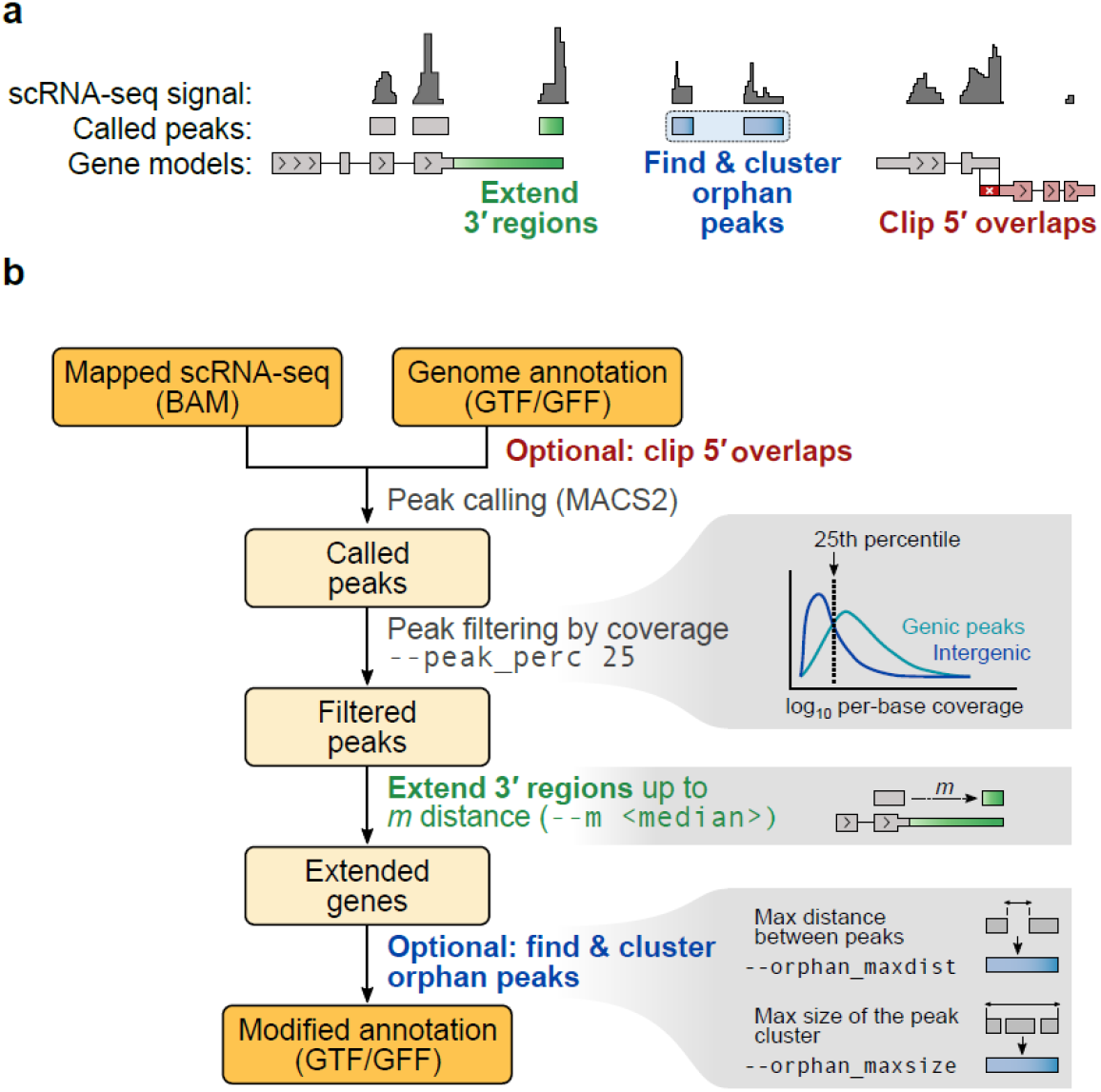
GeneExt tool overview. **a**, GeneExt main function. **n**, Schematic representation of the main steps and associated options.

1. Extension of 3′ regions of known genes to better capture reads from 3′-biased scRNA-seq technologies. This is the core functionality of GeneExt and is performed by default. First, GeneExt will define transcriptionally active regions using a user-provided BAM alignment file (e.g. the one produced by CellRanger or other tools) and MACS2^10^ to identify stranded peaks, which are classified as genic (mapping to previously annotated gene regions) and intergenic. Intergenic peaks are used to extend the 3′ regions of nearby genes in the same strand, up to a certain distance (*--m* flag). Spurious peaks are removed by excluding those with low coverage (by default, intergenic peaks with coverage below the 25th percentile of the genic peak coverage distribution are excluded; *--peak_perc* flag).
2. Identification of orphan peaks that could constitute unannotated genes or longer 3′ UTRs. This is enabled with the *--orphan* flag. Here, GeneExt uses the intergenic peaks not previously assigned to an upstream gene (and passing the coverage filter threshold) to add putative new genes to the final annotation. GeneExt attempts to merge these peaks into clusters defined by a maximum distance between the peaks (-*-orphan_maxdist* parameter, 95th quantile of intron length distribution by default) and filtering out peak clusters above a maximum size (*--orphan_maxsize* parameter, median gene length by default). The idea is to avoid peaks representing exons from the same unannotated gene to contribute independently with the same expression signal in the final UMI count matrix. Clustering of orphan peaks can be disabled with the *--nomerge* flag.
3. Clipping of 5′ regions in cases where they overlap with the 3′ regions of nearby genes. Depending on the behavior of the UMI demultiplexing software used, this overlap can cause (i) the upstream gene to not be quantified (if 3′ biased scRNA-seq reads mapped into the overlapping region are discarded) or (ii) the downstream gene to have two distinct confounding expression signals (if reads are assigned to both). This optional clipping procedure resolves this ambiguity and is enabled by the *--clip_5prime* flag.
4. In addition, GeneExt will also fix non-standard GTF/GFF files provided by the user (e.g. adding gene features if needed), so as to produce outputs compatible with commonly used UMI demultiplexing software (e.g. 10X CellRanger, STARsolo^11^). By default, it will only report the longest isoform of each gene (accordingly extended).

We tested the effects of these gene annotation modification using published whole-organism scRNA-seq atlases from diverse species (**Fig. 2**): the sponge *Amphimedon queenslandica*, the ctenophore *Mnemiopsis leidyi*, the placozoan *Trichoplax adhaerens*^12^, the cnidarians *Nematostella vectensis*^13^, *Stylophora pistillata*^14^ and *Xenia* sp.^15^, the cephalopod *Octopus vulgaris* (in this case only neural tissues)^16^, and the rat *Rattus norvergicus* (nucleus accumbens cells)^17^. Together, they represent not only divergent animal lineages, but also diverse scRNA-seq technologies and different genome assembly and annotation qualities. For each species, we quantified single-cell gene UMI counts for three sets of gene annotations: (*i*) the original annotation, (*ii*) GeneExt-extended gene models, and (*iii*) extended gene models plus orphan peaks. In all cases we used default GeneExt parameters, e.g. to filter out intergenic peaks based on genic peak coverage distributions (**Fig. 2a**).

**Figure 2.**
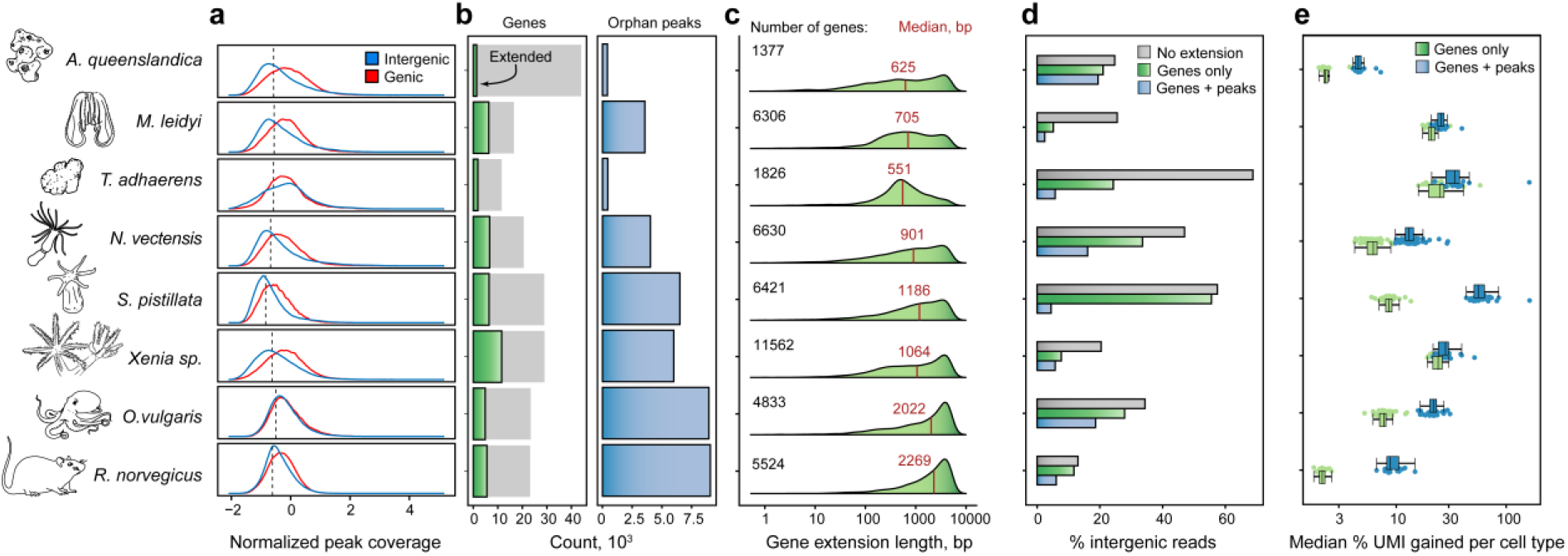
GeneExt application to cell atlases from different species. **a**, Read coverage in called peaks intersecting known exons (“genic peaks”) and intergenic regions. The dashed line indicates the default 25th percentile of genic peaks coverage distribution used to filter intergenic peaks. **b**, Barplots indicating the number of extended genes and number of orphan peaks called in different species using GeneExt default parameters. **c**, Distributions of gene extension lengths across species. The number of extended genes and the median extension length are indicated. **d**, Barplots indicating the percentage of scRNA-seq reads mapping to intergenic regions before and after extending gene annotations in each species. **e**, Boxplots indicating median single-cell UMI gains after extending gene annotations and stratified by cell types (individual dots). Gains are calculated as a median percentage of increase relative to the original UMI counts in each cell.

The fraction of extended genes (**Fig. 2b**) varies greatly across species, from 3.2% in *A. queenslandica* to 39.9% in *Xenia sp*., while the number of remaining intergenic orphan peaks goes from 416 in *A. queenslandica* to 9,079 in *R. norvegicus*. Similar differences can be found in the length of gene extension (**Fig. 2c**). This reflects the fact that incomplete or inaccurate gene models, gene over-annotation and/or under-annotation will affect each species differently. For example, the high fraction of modified genes in *Xenia sp*. probably reflects the systematic mis-annotation of 3′ UTRs in this genome. The different sources of annotation biases are also reflected by observed differences in intergenic mapping reduction after extension (**Fig. 2d**). The high reduction in *T. adhaerens* and *M. leidyi* suggests that most missing information came from 3′-incomplete gene models rather than missing genes. In *S. pistillata*, on the other hand, we observed only a small reduction of intergenic reads after extension, which could be explained by a substantial number of missing genes.

Finally, we tested the gain of UMI counts in each single-cell (**Fig. 2e**) and for each cell type. As expected, inclusion of orphan peaks results in higher UMI gains than simply performing 3′ gene extension. It is apparent that these effects are highly species-specific (again, related with the varying quality of gene annotations in different species), but also cell type-specific, indicating that GeneExt can rescue transcriptomic signal from previously undercounted cell types. This latter effect could be explained by different factors: (*i*) genes that are expressed in rare cell types could be absent in the bulk RNA-seq experiments commonly used for evidence-based gene annotation; or (*ii*) some specialized cell types dedicate a large fraction of their transcriptional output to one or a few genes (e.g. secretory or digestive cells producing proteases), and if these genes belong to families that are systematically mis-annotated for any reason (e.g. they are short or repetitive), this bias will have an outsized effect on these particular cells. Overall, these analyses demonstrate that GeneExt is able to ameliorate the often unanticipated effects that gene annotation inaccuracies can have in the transcriptomes of particular species and cell types.

In summary, GeneExt is a versatile tool to adjust existing gene annotations in order to improve scRNA-seq quantification across species. The software requires minimal input (a pre-existing annotation in any format and scRNA-seq reads) and can be used with default parameters suitable for most species. The result is improved gene detection that facilitates the interpretation of single-cell atlases. Moreover, homogenizing gene detection and quantification in different species is crucial for comparative analysis of cell type gene expression, which is particularly relevant in the context of the taxonomic expansion in cell type atlases across the tree of life.

GeneExt is freely available on Github (https://github.com/sebepedroslab/GeneExt) under a GNU General Public license, together with test data and usage instructions.

## Acknowledgements

Research in A.S-P. group was supported by the European Research Council (ERC-StG 851647) and the Spanish Ministry of Science and Innovation (PID2021-124757NB-I00). We also acknowledge support of the Spanish Ministry of Science and Innovation to the EMBL partnership, the Centro de Excelencia Severo Ochoa and the CERCA Programme (Generalitat de Catalunya). G.Z. is supported INPhINIT PhD fellowship from LaCaixa Foundation LCF/BQ/DI21/11860036. X.G-B. is supported by the European Union’s H2020 research and innovation program under Marie Skłodowska-Curie grant agreement 101031767.

## Methods

### GTF/GFF pre-processing

Genome annotation is usually represented by a tabular file where each row corresponds to a single genomic feature. The hierarchical relationships between features are stored in the 9-th column of the file. Thus, any re-ordering of the file creates problems for the downstream tools that aim to infer such relationships. Another relatively common problem is missing unique IDs for the features. In the case of one-transcript-per-gene annotations, the transcripts are often assigned the same IDs as their parenting genes which makes these IDs non-unique. GeneExt attempts to solve some of these problems by the following:

1. It uses gffutils to parse the hierarchical relationships between the features
2. It adds missing “gene” features
3. It selects a single (longest) transcript per gene
4. It only outputs relevant attributes such as “ID” and “Parent”, in case of GFF or “gene_id”/”transcript_id” in case of GTF

In addition, GeneExt tries to resolve the overlaps between genes on the same strand by:

1. Removing the genes fully contained within another gene
2. Giving priority to the upstream gene

Giving priority to an upstream gene is motivated by the 3′ bias in the single-cell RNA-seq data. A signal is more likely to come from a 3′-UTR of an upstream gene than from the 5′-UTR of the downstream gene.

### BAM file processing and peak calling and filtering

If requested, the alignment file is subsampled (*--subsamplebam*). The reads are then split by strand and peaks are called in each strand using MACS2 software^10^ using the following parameters:

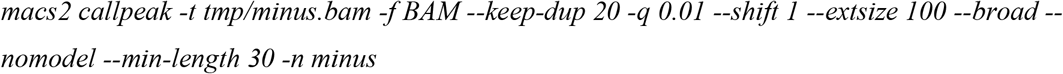

### Gene extension and peak clustering

Called peaks are filtered based on the scRNA-seq coverage, calculated per peak as the total coverage divided by the length of the peak. Then, the coverage of the intergenic peaks is compared to the coverage of the peaks falling within genic regions. Intergenic peaks with normalized coverage exceeding the n-th percentile of genic peak coverage (*--peak_perc*; 25th percentile by default) are then kept for extension.

### Re-analysis of non-model organism single-cell atlases. For each of the species, three versions of the gene annotations were generated

1. Original annotation refined by GeneExt (*--clip_5prime*)
2. Output of GeneExt (*--clip_5prime, -m* 5000, *--subsamplebam* 100000000)
3. Output of GeneExt (*--clip_5prime, -m* 5000, *--subsamplebam* 100000000, *--orphan*)

That is: subsample the dataset to 100M reads, extend the genes to maximum 5000 bp downstream; clip 5′ overlaps in the genome annotation; use 25th coverage percentile for peak filtering; keep orphan peaks.

For single-cell atlases obtained with the MARS-seq scRNA-seq technology (*A. queenslandica, M. leidyi, T. adhaerens, N. vectensis*, and *S. pistillata*), we first mapped reads onto the corresponding genome using STAR 2.7.3^18^, with parameters: *--outFilterMultimapNmax* 20 *---outFilterMismatchNmax* 8 *--alignIntronMax* 3500. Then we quantified gene expression for each of the three interval sets described above using the MARS-seq pipeline as previously described ^19^.

For *Xenia* sp. we only used the tentacle dataset (10x Chromium v2) and for *O. vulgaris* we only used the single-cell RNA-seq dataset (10x Chromium v3), not single-nuclei data. The R2 reads from each dataset were aligned using STAR v2.7.10a^18^ to the corresponding genomes^15,20^. The resulting alignment files were used as an input for GeneExt with the parameters specified above. The resulting genome annotations were used as an input to STARsolo^11^ with default parameters (--*soloCBlen* 16 *--soloUMIlen* 12 was used in the case of *O. vulgaris* to account for v3 Chromium chemistry).

## Notes

https://github.com/sebepedroslab/GeneExt

### Competing Interest Statement

The authors have declared no competing interest.

### Summary of Updates

The manuscript has been revised and updated

## References

1. Tanay, A. & Sebé-Pedrós, A. Evolutionary cell type mapping with single-cell genomics. Trends in Genetics 37, 919–932 (2021).

2. Mudge, J. M. & Harrow, J. The state of play in higher eukaryote gene annotation. Nat Rev Genet 17, 758–772 (2016).

3. Guigó, R. Genome annotation: From human genetics to biodiversity genomics. Cell Genomics 3, 100375 (2023).

4. Amaral, P. et al. The status of the human gene catalogue. Nature 622, 41–47 (2023).

5. Zolotarov, G. et al. MicroRNAs are deeply linked to the emergence of the complex octopus brain. Sci Adv 8, (2022).

6. Legnini, I., Alles, J., Karaiskos, N., Ayoub, S. & Rajewsky, N. FLAM-seq: full-length mRNA sequencing reveals principles of poly(A) tail length control. Nat Methods 16, 879–886 (2019).

7. Wang, M. F. Z. et al. Uncovering transcriptional dark matter via gene annotation independent single-cell RNA sequencing analysis. Nat Commun 12, 2158 (2021).

8. Haese-Hill, W., Crouch, K. & Otto, T. D. peaks2utr: a robust Python tool for the annotation of 3′ UTRs. Bioinformatics 39, 2–3 (2023).

9. Weisman, C. M., Murray, A. W. & Eddy, S. R. Mixing genome annotation methods in a comparative analysis inflates the apparent number of lineage-specific genes. Current Biology 32, 2632-2639.e2 (2022).

10. Zhang, Y. et al. Model-based analysis of ChIP-Seq (MACS). Genome Biol 9, R137 (2008).

11. Kaminow, B., Yunusov, D. & Dobin, A. STARsolo: accurate, fast and versatile mapping/quantification of single-cell and single-nucleus RNA-seq data. bioRxiv 2021.05.05.442755 (2021) doi:10.1101/2021.05.05.442755.

12. Sebé-Pedrós, A. et al. Early metazoan cell type diversity and the evolution of multicellular gene regulation. Nat Ecol Evol 2, 1176–1188 (2018).

13. Sebé-Pedrós, A. et al. Cnidarian Cell Type Diversity and Regulation Revealed by Whole-Organism Single-Cell RNA-Seq. Cell 173, 1520–1534.e20 (2018).

14. Levy, S. et al. A stony coral cell atlas illuminates the molecular and cellular basis of coral symbiosis, calcification, and immunity. Cell 184, 2973-2987.e18 (2021).

15. Hu, M., Zheng, X., Fan, C.-M. & Zheng, Y. Lineage dynamics of the endosymbiotic cell type in the soft coral Xenia. Nature 582, 534–538 (2020).

16. Styfhals, R. et al. Cell type diversity in a developing octopus brain. Nat Commun 13, 7392 (2022).

17. Savell, K. E. et al. A dopamine-induced gene expression signature regulates neuronal function and cocaine response. Sci Adv 6, (2020).

18. Dobin, A. et al. STAR: ultrafast universal RNA-seq aligner. Bioinformatics 29, 15–21 (2013).

19. Keren-Shaul, H. et al. MARS-seq2.0: an experimental and analytical pipeline for indexed sorting combined with single-cell RNA sequencing. Nat Protoc 14, 1841–1862 (2019).

20. Destanović, D. et al. A chromosome-level reference genome for the common octopus, Octopus vulgaris (Cuvier, 1797). G3: Genes, Genomes, Genetics jkad220 (2023) doi:10.1093/g3journal/jkad220.

